# The *Lanata* trichome mutation increases stomatal conductance and reduces leaf temperature in tomato

**DOI:** 10.1101/2020.02.11.943761

**Authors:** Karla Gasparini, Ana Carolina R. Souto, Mateus F. da Silva, Lucas C. Costa, Cássia Regina Fernandes Figueiredo, Samuel C. V. Martins, Lázaro E. P. Peres, Agustin Zsögön

**Affiliations:** Laboratory of Hormonal Control of Plant Development. Departamento de Ciências Biológicas, Escola Superior de Agricultura “Luiz de Queiroz”, Universidade de São Paulo, CP 09, 13418-900, Piracicaba, SP, Brazil; Departamento de Biologia Vegetal, Universidade Federal de Viçosa, CEP 36570-900, Viçosa, MG, Brazil

**Author notes:** **Corresponding author:** Agustin Zsögön, Universidade Federal de Viçosa, Brazil.

**Keywords:** Gas exchange, *Solanum lycopersicum*, stomatal conductance, stress, tomato mutants, transpiration, water loss

## Abstract

**Background and aims:** Trichomes are epidermal structures with an enormous variety of ecological functions and economic applications. Glandular trichomes produce a rich repertoire of secondary metabolites, whereas non-glandular trichomes create a physical barrier against biotic and abiotic stressors. Intense research is underway to understand trichome development and function and enable breeding of more resilient crops. However, little is known on how enhanced trichome density would impinge on leaf photosynthesis, gas exchange and energy balance.

**Methods:** Previous work has compared multiple species differing in trichome density, instead here we analyzed monogenic trichome mutants in a single tomato genetic background (cv. Micro-Tom). We determined growth parameters, leaf spectral properties, gas exchange and leaf temperature in the *hairs absent* (*h*), *Lanata* (*Ln*) and *Woolly* (*Wo*) trichome mutants.

**Key results:** Shoot dry mass, leaf area, leaf spectral properties and cuticular conductance were not affected by the mutations. However, the *Ln* mutant showed increased carbon assimilation (*A*) possibly associated with higher stomatal conductance (*g*_s_), since there were no differences in stomatal density or stomatal index between genotypes. Leaf temperature was furthermore reduced in *Ln* in the early hours of the afternoon.

**Conclusions:** We show that a single monogenic mutation can increase glandular trichome density, a desirable trait for crop breeding, whilst concomitantly improving leaf gas exchange and reducing leaf temperature.

**HIGHLIGHT:** *A monogenic mutation in tomato increases trichome density and optimizes gas exchange and leaf temperature*

## INTRODUCTION

Trichomes are extensions of the epidermis found in most terrestrial plants. The density and type of trichomes are under strong genetic control but can vary due to environmental factors (Holeski *et al.*, 2010; Thitz *et al.*, 2017). Variation is inter- and intra-specific, but may occur within an individual as well, depending on leaf position and developmental stage of the plant (Chien and Sussex, 1995; Vendemiatti *et al.*, 2017). Glandular trichomes are characterized by the ability to synthesize, store and secrete a vast array of secondary metabolites, mostly terpenoids and phenylpropanoids, making them appealing platforms for biotechnological applications (Tissier, 2012; Huchelmann *et al.*, 2017). Furthermore, considerable interest has arisen in exploiting glandular trichomes as a breeding tool to increase resistance to herbivory and thus avoid excessive use of pesticides (Simmons and Gurr, 2005; Maluf *et al.*, 2007; Tian *et al.*, 2014; Chen *et al.*, 2018). Thus, research has focused on identifying metabolic pathways within glandular trichomes (Besser *et al.*, 2009; Sallaud *et al.*, 2009; Bleeker *et al.*, 2012; Balcke *et al.*, 2017; Leong *et al.*, 2019).

However, some important gaps in the knowledge need to be resolved before increased trichome density can be deployed in crops. Leaf pubescence caused by high trichome density can have direct and indirect effects on photosynthesis, transpiration and leaf energy balance (Ehleringer and Mooney, 1978; Bickford, 2016). On the one hand, modelling work has shown that trichomes may affect gas exchange through increasing the leaf boundary layer resistance (Schreuder *et al.*, 2001). The boundary layer is a zone of relatively calm air that can influence the transfer of heat, CO_2_ and water vapour between the leaf and the environment (Schuepp, 1993). Research, however, has shown that this effect is only significant under particular environmental conditions (Ehleringer and Mooney, 1978; Benz and Martin, 2006). On the other hand, a more pronounced effect is the increased reflectance of photosynthetically active radiation (PAR) by highly pubescent leaves, which can occur by trichomes themselves (Sandquist and Ehleringer, 1997, 1998) or by accumulation of water droplets (Brewer *et al.*, 1991). Altered light absorptance can in turn affect photosynthetic rate directly through decreased PAR or indirectly through increased leaf temperature (Bickford, 2016).

Tomato (*Solanum lycopersicum*) and its related wild species display great diversity in density and type of trichomes: seven types of trichome, of which three (types II, III and V) are non-glandular and four (types I, IV, VI and VII) correspond to the glandular type (Luckwill, 1943; Glas *et al.*, 2012). Tomato has, therefore, been proposed as a model for the study of trichomes (Schuurink and Tissier, 2019). Here, we investigated how changes in trichome density influence leaf spectral properties, gas exchange and leaf temperature. We used four tomato (cv. Micro-Tom, MT) genotypes that differ in trichome density: wild-type (MT) and mutants with either decreased (*hairs absent, h*), or increased (*Lanata, Ln* and *Woolly, Wo*) trichome density. We found that the *Ln* mutation increased assimilation rate and stomatal conductance, while reducing leaf temperature. Given the growing interest in altering trichome density in tomato and other crops to increase insect resistance (Glas *et al.*, 2012), and to exploit the production of secondary metabolites (Sridhar *et al.*, 2019), we discuss the physiological and breeding implications of our results.

## MATERIAL AND METHODS

### Plant material and experimental setup

Seeds of tomato (*Solanum lycopersicum* L) mutants in their original background were donated by Dr. Roger Chetelat (Tomato Genetics Resource Center – TGRC, Davis, University of California) and then introgressed into the Micro-Tom (MT) genetic background to generate near-isogenic lines (NILs). The NILs harbouring the mutations *Lanata* (*La*), *Woolly* (*Wo*), *hair absent* (*h*) were generated as described by (Carvalho *et al.*, 2011). Seeds of MT were kindly donated by Prof. Avram Levy (Weizmann Institute of Science, Israel) in 1998 and kept as a true-to-type cultivar through self-pollination. All the steps in the introgression process were performed in the Laboratory of Hormonal Control of Plant Development (HCPD), Escola Superior de Agricultura “Luiz de Queiroz” (ESALQ), University of São Paulo, Brazil.

Plants were grown in a greenhouse at the Federal University of Viçosa (642 m asl, 20° 45’ S; 42° 51’ W), Minas Gerais, Brazil, under semi-controlled conditions. Seeds were germinated in 2.5 L pots containing commercial substrate (Tropstrato® HT Hortaliças, Vida Verde). Upon appearance of the first true leaf, seedlings of each genotype were transplanted to pots containing commercial substrate supplemented with 8 g L^−1^ 4:14:8 NPK and 4 g L^−1^ dolomite limestone (MgCO_3_ + CaCO_3_).

### Biomass measurement

Shoot and root biomass were measured from the dry weight of leaves, stem and roots after oven-drying at 80 °C for 24 h. The measurement was performed in plants 40 days after germination. Leaf area was measured by digital image analysis using a scanner (Hewlett Packard Scanjet G2410, Palo Alto, California, USA), and the images were later processed using the ImageJ (NIH, Bethesda, Maryland, USA).

### Leaf spectral properties

Spectral analysis of leaves was performed in the attached central leaflet of the fourth fully expanded leaf 56 days after germination. Leaf reflectance and transmittance were measured on the adaxial side of the leaflet throughout the 280–880 nm spectrum using a Jaz Modular Optical Sensing Suite portable spectrometer (Ocean Optics, Inc., Dunedin, Florida, USA).

### Gas exchange

The net rate of carbon assimilation (*A*), stomatal conductance (*g*_s_), internal CO_2_ concentration (C_i_) and transpiration (*E*) were measured in the third or fourth (from the bottom) fully expanded leaf in all genotypes 40 days after germination. Gas exchange measurements were performed using an open-flow gas exchange infrared gas analyzer (IRGA) model LI-6400XT (LI-Cor, Lincoln, NE, USA). The analyses were performed under common conditions for photon flux density (1000 μmol m^−2^ s^−1^, from standard LiCor LED source), leaf temperature (25 ± 0.5°C), leaf-to-air vapor pressure difference (16.0 ± 3.0 mbar), air flow rate into the chamber (500 μmol s^−1^) and reference CO_2_ concentration of 400 ppm (injected from a cartridge), using an area of 6 cm^2^ in the leaf chamber.

### Epidermal features analysis

Dental resin impressions were used to obtain number and area of pavement cells and stomatal density, following the methodology described by (Geisler *et al.*, 2000). The third or fourth (from the bottom) fully expanded leaf was sampled 56 days after germination for resin impressions in all genotypes. Using a spatula, the mixture of Oranwash L and indurent gel (Zhermack) was applied to the adaxial and abaxial face of opposite leaflets. After drying, the resin mold was removed from the leaflet surface and filled with nail polish to create a cast that was analyzed by light microscopy (Zeiss Axioskop 40, Carl Zeiss, Jena, Germany). Images were analyzed with Image-Pro ® PLUS. Stomatal index (SI) was calculated using the formula SI=S/(S+E) ×100, where S is number of stomata per area and E is number of epidermal cells per area. The analysis was performed in five plants per genotype, five images per plant, totalizing 25 images per genotype. Pavement cell area were measured for 30 pavement cells per genotype.

### Leaf temperature determinations

Leaf temperature was measured in the central leaflet of the fourth fully expanded leaf 60 days after germination using an infrared thermometer (LASERGRIP GM400). Measurements were performed from 8:00 am to 6:00 pm with an interval of two hours between each measurement. A second set of measurements was carried out on thermal images obtained simultaneously with an infrared camera (FLIR systems T360, Nashua USA). All image processing and analysis was undertaken in Flir Tools software version 5.2.

### Minimum epidermal conductance (g_mim_)

Analysis of minimum epidermal conductance was performed in plants 62 days after germination. The fully expanded fourth leaf was carefully removed and the petiole was sealed with silicone. The leaves were placed in zip lock bags with high CO_2_ concentration. After an hour in the dark, the leaves were weighed on an analytical balance every 15 minutes for four hours. At the end of measurements, leaf area was calculated digitizing leaves with an HP Scanjet G2410 scanner (Hewlett-Packard, Palo Alto, California, USA) and performing image analysis with ImageJ. The minimum cuticular conductance was calculated using the *g*_min_ Analysis Worksheet Tool Created by Lawren Sack (University of California, Los Angeles, http://prometheuswiki.org).

### Transpirational water loss under still and turbulent air

Transpirational water loss determinations were performed in 56-day old plants. In order to evaluate the influence of the boundary layer on transpiration, measurements were performed under either static or turbulent air conditions. Turbulent air was created with a fan (ARNO, São Paulo, Brazil) placed at 1.5m distance, at the minimum ventilation velocity (determined to be 10 km/h using an anemometer, Bmax, Anemometer). We determined the ‘static’ air to be 1 km/h using the same instrument. Pots were irrigated the night before the measurements until they reached field capacity. Later the pots were covered with plastic film to reduce water loss from the substrate via evaporation and then weighed. The following day, pots were weighed every two hours between 8:00 am and 6:00 pm. Water loss was calculated as the difference in weight every between measurements.

### Statistical analyses

The data were obtained from the experiments using a completely randomized design with the genotypes MT, *h, Ln* and *Wo*. Data are expressed as mean ± s.e.m. Data were subjected to one-way analysis of variance (ANOVA), followed by Tukey’s honestly significant difference test (*p*<0.05) or Fisher’s Least Significance Difference (LSD) test. All statistical analyses were carried out using Assistat v. 7.7 (Campina Grande, Brazil).

## RESULTS

### Mutations altering trichome type and density do not affect leaf area or shoot dry mass

Here, we studied three near-isogenic lines (NILs) that harbor mutations affect trichome density in the genetic background of tomato cv. Micro-Tom (MT) (see Supplementary Table 1 for details). Like most cultivated tomatoes, MT displays three types of glandular trichomes (types I, VI and VII) and two types of non-glandular trichomes (III and IV) in adult leaves. Typically, total trichome density higher on the abaxial (around 250 trichomes cm^−2^) than on the adaxial (∼100 trichomes cm^−2^) side of the leaf, and the most abundant on both sides is type V (Fig. 1). The *h* mutant displays a small, but significant reduction on type III and VI trichome density. Both the *Ln* and *Wo* mutants show higher trichome density on both sides of the leaf. The change, however, is not proportional among trichomes types: both *Ln* and *Wo* show preferential increases of types III and V (adaxial) and I and V (abaxial). Both genotypes also display a branched (‘ramose’) type of trichome that is absent in MT. Altogether, these changes result in a very similar ‘hairy leaf’ phenotype for *Ln* and *Wo*, with roughly three-fold increase of total trichome density on either side of the leaf.

**Figure 1.**
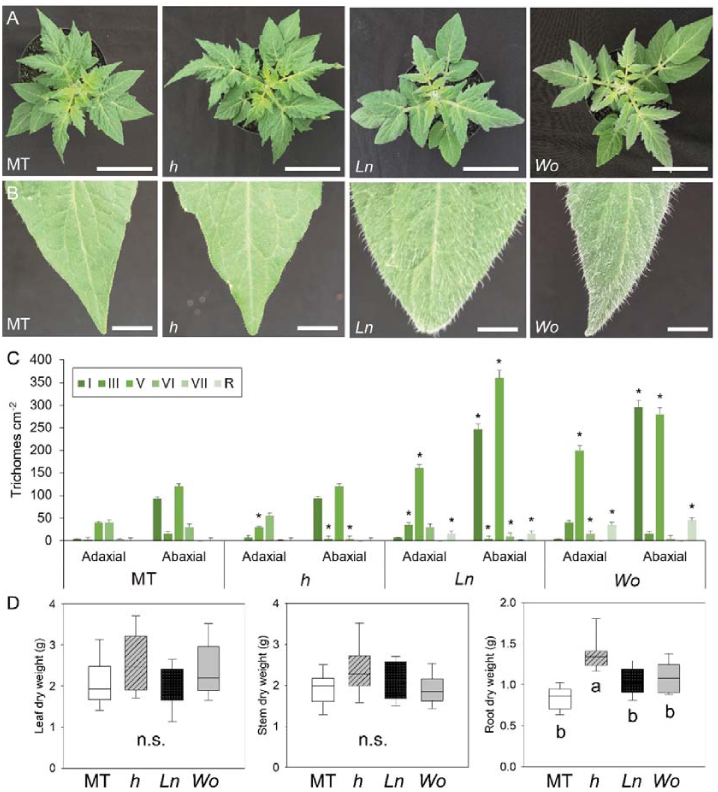
Macroscopic view of representative plants of tomato cv. Micro-Tom (MT) and the trichome mutants *hairs absent* (*h*), *Lanata* (*Ln*), *Woolly* (*Wo*). (A) Top view, bar = 5 cm. (B) Detail of leaf tips showing the difference in pilosity between genotypes, bar = 1cm. (C) Trichome density per trichome type on the adaxial and abaxial side of fully-expanded leaves (based on Vendemiatti et al. 2017). Asterisks indicate significant differences to MT (*p*<0.05). (D) Leaf, stem and roots dry weight determined through destructive analysis in plants 65 days after germination. Values are means ± s.e.m. (n=10). Significant differences tested with one-way ANOVA followed by Tukey’s honestly significant difference (HSD) test, letters indicate significant differences (*p*<0.05). n.s. indicates that no significant difference was found between genotypes.

Given the possibility of unrelated pleiotropic alterations caused by the mutations, we assessed other macroscopic plants traits. Except for a subtle reduction in leaf margin indentation in *Ln* (Fig. 1), we found no differences in leaf architecture, leaf area or shoot dry mass between the mutants of trichomes and MT (Fig. 1). Remarkably, all the *h* mutant showed higher root system dry mass (Fig. 1).

### Changes in trichome density do not affect the leaf spectral properties

Depending on their density, trichomes may affect leaf function by altering the amount of light absorbed by leaves and/or by increasing the resistance of the boundary layer. Thus, we next measured transmittance and reflectance of photosynthetically active radiation (PAR) on leaves of MT and the trichome mutants. The *h* mutant had a lower reflectance than the *Ln* mutant (*p*=0.0068), but neither was significantly different to MT or to the *Wo* mutant. No differences were found in leaf transmittance or absorptance in the PAR wavelength (Fig. 2). Calculation of the boundary layer resistance is problematic (see Discussion). To provide an indirect assessment of the effect of trichomes on the boundary layer, we next carried out a gravimetric determination of transpirational water loss under conditions of static (1 km h^−1^) and turbulent (10 km h^−1^) air. Under static air, we did not observe differences in rates of water loss between genotypes throughout the day or in transpiration calculated as water lost divided by leaf area over time (Fig. S1). Under turbulent air, water loss was lower for all genotypes, but without differences between them, except for one interval (between 12:00-14:00) for the *Ln* mutant, which showed less water loss than MT (Fig. S1).

**Figure 2.**
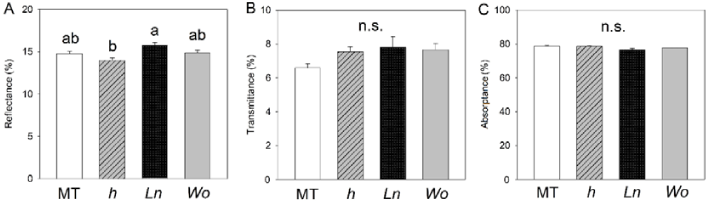
Leaf spectral properties are not affected by trichome density in tomato. Percentage reflectance (A), transmittance (B) and absorptance (C) on leaves of Micro-Tom (MT) and the *hairs absent* (*h*), *Lanata* (*Ln*) and *Woolly* (*Wo*) mutants, measured 56 days after germination. Values are means ± s.e.m. (n=6). Significant differences tested with one-way ANOVA followed by Tukey’s honestly significant difference (HSD) test, letters indicate significant differences (*p*<0.05). n.s. indicates that no significant difference was found between genotypes.

### *The* Ln *mutation increases photosynthetic assimilation rate and stomatal conductance*

We next investigated the effects of trichome mutations on leaf gas exchange. Both net photosynthesis rate (*A*) and stomatal conductance (*g*_s_) were 29% and 38% higher in *Ln* than in MT (*p*<0.0001), respectively (Fig. 3). Internal CO_2_ concentration (*C*_i_) was higher in *h* (12%) and *Wo* (9.5%) mutants, compared with MT, but we found no difference in the *C*_i_ of *Ln* mutant (Fig. 3). Intrinsic water-use efficiency (*A/g*_s_) was higher in MT and *Ln* than in *h* and *Wo*.

**Figure 3.**
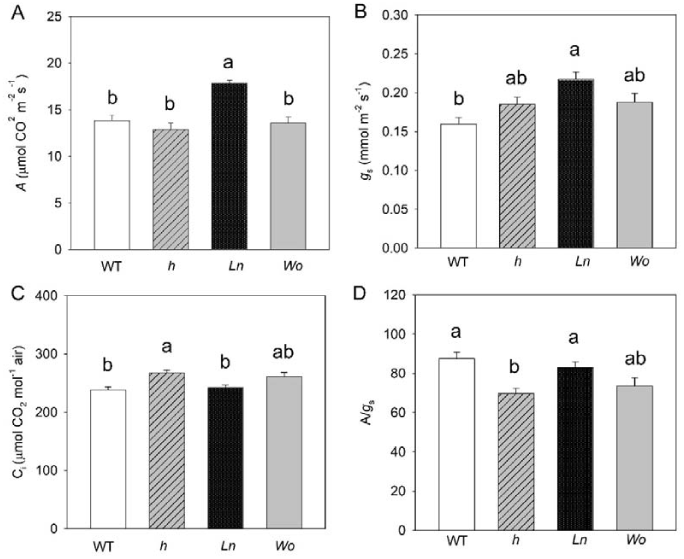
Trichome mutants affect leaf gas exchange parameters in tomato. Measurements were performed on fully expanded leaves of Micro-Tom (MT) and the *hairs absent* (*h*), *Lanata* (*Ln*) and *Woolly* (*Wo*) mutants 40 days after germination. (A) Net carbon assimilation rate (*A*). (B) Stomatal conductance (*g*_s_). (C) Internal CO_2_ concentration (*C*_i_). (D) Intrinsic water-use efficiency (*A*/*g*_s_) calculated using the data from panels A and B. Values are mean ± s.e.m. (n=8). Significant differences tested with one-way ANOVA followed by Tukey’s honestly significant difference (HSD) test, letters indicate significant differences (*p*<0.05).

Given the changes in *g*_s_, we determined whether trichome mutations have an impact on stomatal size and density. Stomatal density can be altered by changes on pavement cell size and density. Pavement cell density was higher in *h* and *Ln* than in *MT* and *Wo* on the adaxial side of the leaf (Fig. S2). Pavement cell area was higher in MT than in all three trichome mutants. However, neither stomatal density nor stomatal index were different between genotypes (Fig. S2). Stomatal size can influence conductance, so we also determined the length and width of the guard cells. Except for a significant guard cell length reduction in *Ln* (restricted to the adaxial side of the leaf), no differences were found in stomatal size between genotypes (Fig. S3).

Stomata and cuticle are the primary leaf components responsible for maintaining plant water status (Schuster *et al.*, 2016). To avoid excessive water loss, stomata close in periods of water deficit. Even when stomata are closed, water loss continues at very low rates through the leaf cuticle. This basal rate of water loss is expressed as the minimum conductance of a leaf (*g*_min_). Both *g*_min_ and stomatal closure can be affected by trichomes (Hegebarth *et al.*, 2016; Galdon-Armero *et al.*, 2018). However, we found no differences in *g*_min_ among the genotypes evaluated in this experiment (Supplementary data Fig. S4).

### The *Ln* mutation reduces leaf temperature during the afternoon

Trichome density may potentially influence leaf temperature. To explore this effect, we determined a time course of temperature using two variants of infrared thermography (camera and thermometer). We found a steep increase in leaf temperature over the course of the morning, reaching more than 30°C around noon (Fig. 4). From then on, a further, yet smoother increase occurred until 15:00, when leaf temperature reached 35°C. This, however, was not the case for *Ln* plants, which maintained a lower temperature over the transition from noon to the mid-afternoon. Eventually, leaves of all plants showed a steep decline in temperature at 16:30 and later (Fig. 4).

**Figure 4.**
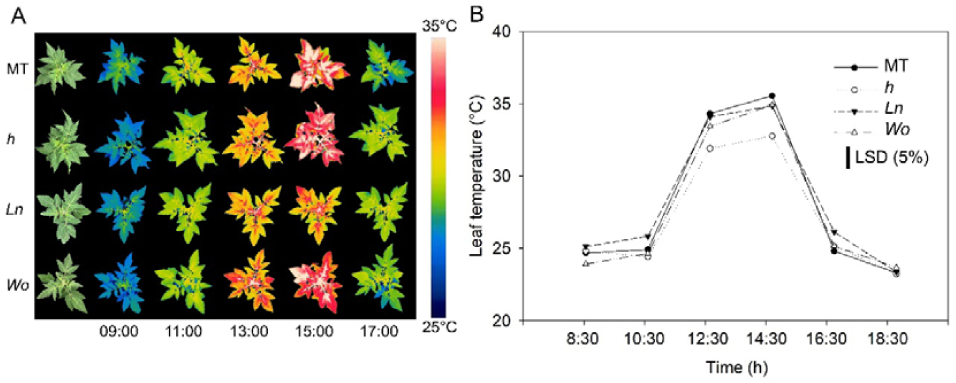
The *Lanata* (*Ln*) mutation reduces afternoon leaf temperature. Time courses of leaf temperature on tomato cv. Micro-Tom (MT) and *hairs absent* (*h*), *Lanata* (*Ln*) and *Woolly* (*Wo*) trichome mutants using (A) infrared thermography (representative plants shown), and (B) point-and-shoot infrared thermometer. Measurements were conducted on fully expanded leaves of 60-day old plants (n=8). LSD: Fisher’s Least Significant Difference between means.

## DISCUSSION

The recent years have seen a growing interest in harnessing trichomes as either a biotechnological platform for the production of valuable metabolites (Huchelmann *et al.*, 2017) or as an alternative, natural source of resistance to multiple biotic and abiotic stressors in crops (Glas *et al.*, 2012; Oksanen, 2018). However, some gaps in basic knowledge need to be resolved before trichome manipulation can be deployed as a breeding tool (Schuurink and Tissier, 2019; Chalvin *et al.*, 2020). Among others, an important question refers to the ecophysiological trade-off between advantages and disadvantages of increasing trichome density in crops (Bickford, 2016). Here we set out to explore this question using monogenic trichome mutants of tomato. Considerable variation in trichome type and density exists in cultivated tomato and its wild relatives, making this a suitable system for trichome study (Simmons and Gurr, 2005; Glas *et al.*, 2012). Monogenic mutants have been described controlling alterations in trichome type and density (Yang *et al.*, 2011*a*,*b*; Chang *et al.*, 2018). Given the epistatic effects of genetic background, functional comparative studies of mutants are best performed using the same tomato cultivar (Tonsor and Koornneef, 2005).

Here, we analyzed trichome mutants in the same tomato cultivar (MT), thus avoiding the confounding effects of variation in genetic background (Carvalho *et al.*, 2011). The only macroscopic alteration we observed was an increase in root dry weight in the *h* mutant. Although inconsistent, changes in root structure are not unusual in trichome mutants (Yang *et al.*, 2011*a*; Khosla *et al.*, 2014; Li *et al.*, 2016). In tomato plants, the expression pattern of the *Woolly* (*Wo*) gene in roots highlights a role for *Wo* in lateral root differentiation (Yang *et al.*, 2011*a*). The absence of leaf trichomes in the *tril* mutant in cucumber was accompanied by short root hairs and increases in root length and branching (Li *et al.*, 2016). On the other hand, *gl2* mutants of *Arabidopsis* exhibit glabrous leaves and excess root hair formation in primary roots (Khosla *et al.*, 2014). Since many genes that control trichome development play a pivotal role in regulating the differentiation of the epidermis in various tissues, including roots, changes in root growth and structure are not unexpected.

Leaf trichomes have long been known to affect light absorptance, which in turn can influence photosynthetic rate and leaf temperature (Ehleringer *et al.*, 1976; Ehleringer and Björkman, 1978). We only found a minor alteration in reflectance of photosynthetically active radiation (PAR) between genotypes, a 1% increase in *Ln* and a >1% reduction in *h* compared to MT. This is probably because even in the pubescent mutants, trichome morphology cannot effect changes on reflectance. A similar result was found comparing pubescent and glabrous populations of another solanaceous species, *Datura wrightii* (Ii and Hare, 2004). On the other hand, closely related species of *Pachycladon* with comparable trichome densities exhibited considerably different spectral properties – an effect that was ascribed to variation in trichome morphology (Mershon *et al.*, 2015).

Trichomes can indirectly affect gas exchange by altering light absorptance, boundary layer resistance and/or leaf temperature. Determination of the boundary layer resistance is challenging, given the large number of biophysical variables involved (Schuepp, 1993). The current consensus is that the effect of trichomes is negligible compared to stomatal resistance (Bickford, 2016). Comparing transpirational water loss under conditions of still and turbulent air we could find no difference between genotypes, except for slight decrease of water loss in *Ln* under turbulent conditions. Our finding of increased *A* and *g*_s_ in the *Ln* mutant but not in *Wo* was surprising, given the strong resemblance between both phenotypes. The molecular identity of the *Ln* mutation is hitherto unknown, so this effect remains enigmatic. Also, It has been proposed that trichomes could act as dew or fog collectors, wetting the leaf surface and reducing the leaf-to-air water potential difference and thus increasing WUE (Brewer *et al.*, 1991; Brewer and Smith, 1994). Our results do not support this notion, as both *Ln* and *Wo* have similar WUE to the MT control. We could furthermore not ascribe the increased *g*_s_ in *Ln* to alterations on stomatal density or size (in fact, a slight decrease in adaxial guard cell length for the mutant). Thus, it seems plausible that *Ln* is a direct molecular regulator of stomatal opening.

Lastly, leaf temperature was lower in *Ln* during the hot hours of midday and the early afternoon. This effect was not observed in the *Wo* mutant, nor was increased temperature found for *h*, so it does not appear to be a consequence of changes in trichome density, but rather the result of evaporative cooling through increased transpiration caused by a higher *g*_s_. Reduced leaf temperature in *Ln* could have a synergistic positive effect along with increased *g*_s_ on carbon assimilation rate (Ripley *et al.*, 1999). The temperature dependence of C_3_ photosynthesis is quite steep below 20°C and above 30°C, with the optimum around 25 °C (Sage and Kubien, 2007). The reduction of more than 2°C in *Ln*, especially at the hottest hours of the day, could have a beneficial effect for field-grown tomatoes in tropical regions by potentially reducing photoinhibition.

Given the relationship between trichomes and water relations, recent research has evaluated the role of trichomes in tolerance and adaptation to water stress (Sletvold, 2011; Ewas *et al.*, 2016; Mo *et al.*, 2016; Galdon-Armero *et al.*, 2018). In *Arabidopsis lyrata*, plants with higher density of trichomes were more tolerant to drought than glabrous plants (Sletvold, 2011). In watermelon (*Citrullus lanatus*), tolerance to drought stress in wild genotypes, in relation to domesticated ones, is linked to the increased density of trichomes in the former (Mo *et al.*, 2016). In tomato, overexpression of MIXTA-like MYB transcription factor led to higher density of trichomes and tolerance to water stress (Ewas *et al.*, 2016). Further work demonstrated that the ratio of trichomes to stomata plays a role in avoiding excessive water loss (Galdon-armero *et al.*, 2018). Collectively, these results make increasing glandular trichome density in crops an appealing breeding prospect, for instance glandular trichome type IV, which is found in wild tomato relatives but not in adult leaves of cultivated tomato (Vendemiatti *et al.*, 2017).

## CONCLUSIONS

Leaf trichomes are key structures regulating plant-environment interactions. The use of glandular trichomes as a biotechnological platform or as a breeding tool requires a deeper understanding of both the molecular control of their development and the ecophysiological consequences of increased leaf pubescence. We have shown that the monogenic *Lanata* mutation leads to higher assimilation rate and stomatal conductance, resulting in lower leaf temperature, thus bypassing some of the undesirable physiological traits usually associated with leaf pubescence for agronomic settings. Cloning and molecular characterization of *Ln* should pave the way for its exploitation in crop breeding.

## SUPPLEMENTARY DATA

Table S1. Description of the genotypes studied in this work.

Table S2. Leaf area of the plants

Figure S1. Transpirational water loss time-courses

Figure S2. Leaf epidermis features

Figure S3. Guard cell size

Figure S4. Mininum leaf conductance (*g*_min_)

## ACKNOWLEDGEMENTS

This study was financed in part by the Coordination for the Improvement of Higher Level Personnel (CAPES-Brazil) (Finance Code 001).

