## Supplemental Data for "The *Lanata* trichome mutation increases stomatal conductance and reduces leaf temperature in tomato"

**SUPPLEMENTARY DATA**

**Table S1**. Description of tomato (*Solanum lycopersicum* L.) cv Micro-Tom (MT) trichome mutants used in this work. Adapted from (Vendemiatti *et al.*, 2017).

| Genotype | Description | Accession | References |
| --- | --- | --- | --- |
| *hairs absent* (*h*) | Loss-of-function mutant in a C2H2 Zinc Finger Protein (ZFP), related to the *Arabidopsis* ZFP8 gene.  Gene ID: Solyc10g078970. | Introgressed from LA1221 cv. VFNT Cherry | (Chang *et al.*, 2018). |
| *Lanata* (*Ln*) | Mutant with unknown gene function. | Introgressed from LA3128 (*G* allele) | - |
| *Woolly* (*Wo*) | Loss-of-function mutant in an HD-Zip protein related to the *Arabidopsis*’ *PROTODERMAL FACTOR2 (PDF2)* and to *GLABRA2 (GL2)*  mutants, controlling the B type cyclin SlCyCB2.  Gene ID: Solyc02g080260. | Introgressed from LA0715 (*m* allele). | (Yang *et al.*, 2011*a*,*b*). |

**Table S2.** Leaf area determined in plants 62 days after germination. Micro-Tom (MT), *hairs absent* (*h*), *Lanata* (*Ln*), *Woolly* (*Wo*).

| Genotype | Leaf area (cm^2^) |
| --- | --- |
| MT | 449.65 ± 35.48 |
| *h* | 430.02 ± 56.70 |
| *Ln* | 389.18 ± 40.40 |
| *Wo* | 487.11 ± 14.68 |

Values are means ± SEM (n = 4). No significant difference was found with one-way ANOVA.

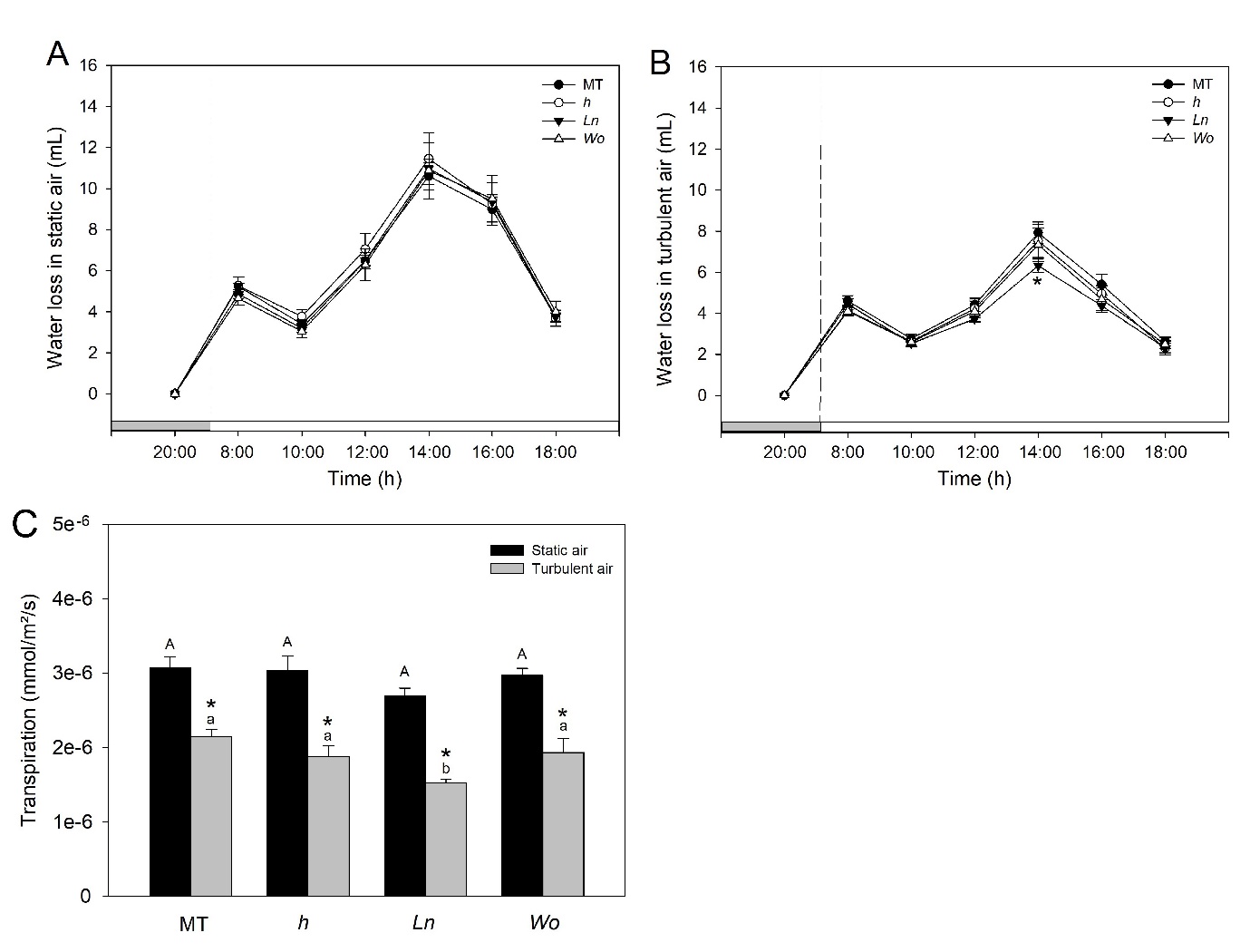

**Figure S1.** Gravimetric transpiration in intact 56-day old plants. (A-B) The grey bar on the x-axis indicates a measurement performed the night when the plant pots were at field capacity. The white bar indicate measurement performed at light period. (C) Bar charts showing accumulated water lost through transpiration derived from plots A and B. Values are means ± s.e.m. (n=6). Significant differences tested with one-way ANOVA followed by Tukey’s honestly significant difference (HSD) test, the asterisks indicates significant difference (*p*<0.05). Capital letters indicate significant differences between genotypes and lower case letters between treatments (*p*<0.05).

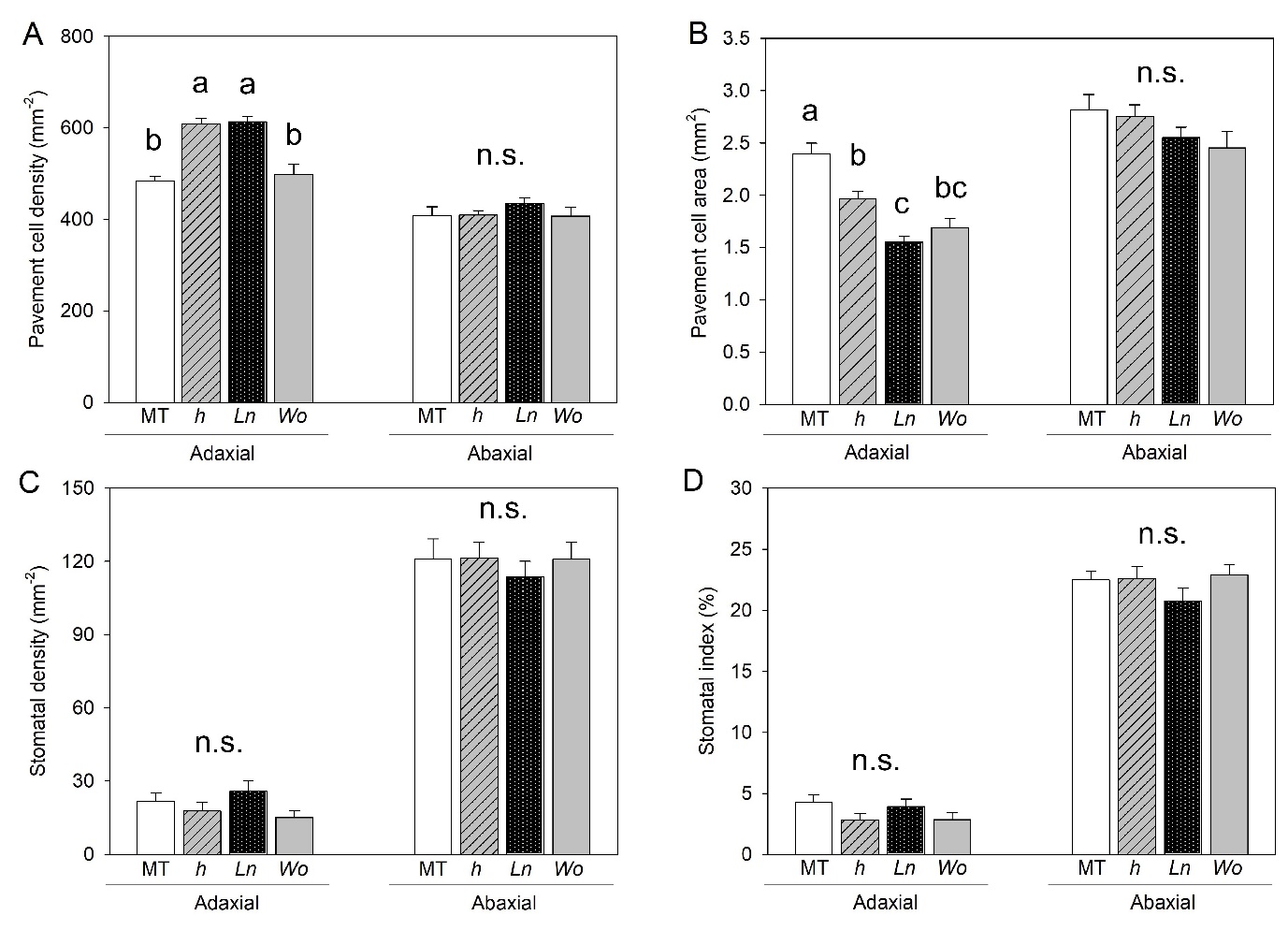

**Figure S2.** Leaf epidermis features in plants of tomato cv. Micro-Tom (MT) and the trichome mutants *hairs absent* (*h*), *Lanata* (*Ln*), *Woolly* (*Wo*). Increase in density of pavement cells on the adaxial leaf face of mutants *h* and *Ln* is due to the reduction in pavement cell area. A, pavement cell density (mm^-2^). B, stomatal density (mm^-2^). C, pavement cell area (mm^2^). D, stomatal index (%). Analyses were performed in plants 55 days after germination. Values are means ± s.e.m. (n=8). Significant differences tested with one-way ANOVA followed by Tukey’s honestly significant difference (HSD) test, letters indicate significant differences (*p*<0.05). n.s. indicates that no significant difference was found between genotypes.

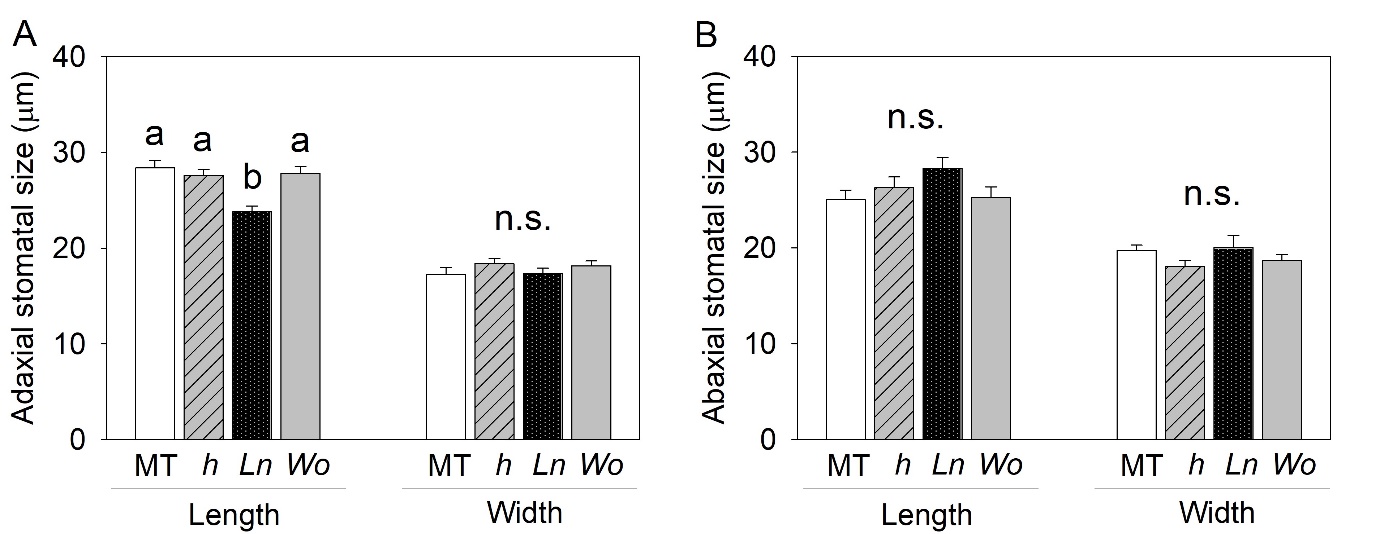

**Figure S3.**  Guard cell length is reduced in the *Lanata* (*Ln*) mutant compared to cv. Micro-Tom (MT) and the trichome mutants *hairs absent* (*h*) and *Woolly* (*Wo*). Values are means ± s.e.m. (n=20). Significant differences tested with one-way ANOVA followed by Tukey’s honestly significant difference (HSD) test, letters indicate significant differences (*p*<0.05).

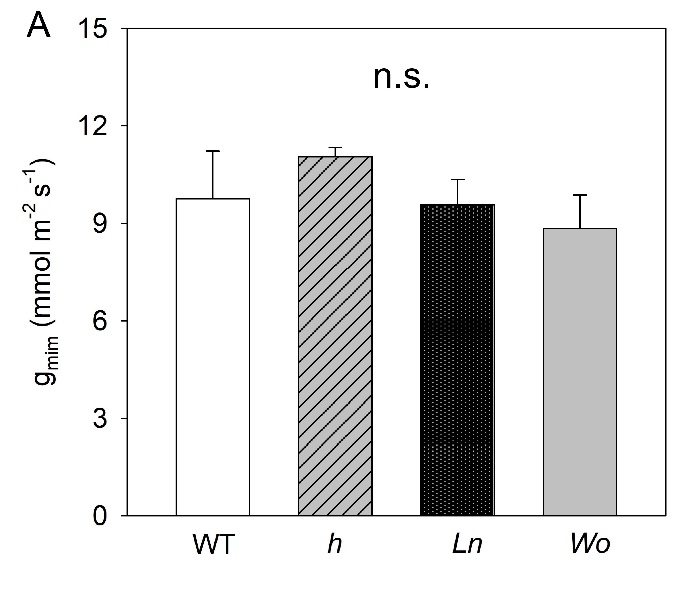

**Figure S4.** Leaf minimum conductance (*g*_min_) determined on fully expanded leaves 62 days after germination. Values are means ± s.e.m. (n=6). Significant differences tested with one-way ANOVA followed by Tukey’s honestly significant difference (HSD) test, n.s. indicates that no significant difference was found between genotypes.
